# Tissue recoil in the early Drosophila embryo is a passive not active process

**DOI:** 10.1101/2022.08.29.505741

**Authors:** Amanda Nicole Goldner, Salena M. Fessehaye, Kelly Ann Mapes, Miriam Osterfield, Konstantin Doubrovinski

## Abstract

Understanding tissue morphogenesis is impossible without knowing the mechanical properties of the tissue being shaped. Although techniques for measuring tissue material properties are continually being developed, methods for determining how individual proteins contribute to mechanical properties are very limited. Here, we developed two complementary techniques for the acute inactivation of sqh (the *Drosophila* myosin regulatory light chain), one based on the recently introduced AID2 system, and the other based on a novel method for conditional protein aggregation that results in nearly instantaneous protein inactivation. Combining these techniques with rheological measurements, we show that passive material properties of the cellularization-stage *Drosophila* embryo are essentially unaffected by myosin activity. These results establish that this tissue is elastic, not predominantly viscous, on the developmentally relevant timescale.

**Summary:** Techniques to examine the contribution of specific proteins to tissue mechanical properties are extremely limited. Here, Goldner et al. develop two complementary techniques for rapid protein depletion combined with mechanical measurements, and show that myosin activity is dispensable for tissue elasticity.

## Introduction

Tissue morphogenesis is at its core a mechanical process, in which changes in tissue shape are directed by spatial-temporal patterns of forces and material properties (Davidson, 2017, Keller, 2012). How the material properties of a tissue - such as elasticity, viscosity, or plasticity - arise from its constituent parts is an area of ongoing investigation. In some tissues, mechanical properties may be dominated by supracellular structures based on extracellular matrix, cell-cell interactions, or cell-interstitial packing (Lenne and Trivedi, 2022). In other tissues, such as simple epithelia with strong cell adhesion, the mechanical properties of cells, and therefore the cytoskeleton, might be expected to dominate tissue properties.

One approach to studying cytoskeletal mechanics involves studies of cytoskeletal gels that have been reconstituted in vitro (Gardel et al., 2008, Joanny and Prost, 2009, Pegoraro et al., 2017). Rheological measurements, combined with theoretical or computational approaches, have generated insights into the mechanical effects of filament length distribution (Janmey et al., 1994), and the effects of adding crosslinkers or motors (Bendix et al., 2008, Koenderink et al., 2009). However, it can be difficult to directly translate these insights into understanding how cells actually control cytoskeletal mechanics in vivo. This is partially because reconstituted systems generally have fewer distinct components that of in vivo cytoskeletal networks; for example, the number of distinct actin binding proteins in the fly has been estimated to be around 80 (Goldstein and Gunawardena, 2000). Additionally, in vitro gels are usually spatially homogenous, while cytoskeletal networks in cells usually are non-homogenous (for example, the cortex versus the bulk of the cytosol) (Van Citters et al., 2006), and can sometimes exhibit exquisite spatial patterning (Dubey et al., 2020, Xu et al., 2013).

A complementary approach has been to conduct mechanical measurements directly on living cells or tissues (Khalilgharibi et al., 2019, D’Angelo et al., 2019, Campas et al., 2014, Mongera et al., 2018, Serwane et al., 2017). These studies are extremely valuable for determining the actual material properties of these complex biological entities. However, studies aimed at understanding the contribution of specific proteins to cell or tissue rheology are much more limited, and have mostly been done with proteins that can be targeted pharmacologically (D’Angelo et al., 2019, Van Citters et al., 2006). In order to more generally assess the role specific proteins play in tissue mechanics, it is critical to measure tissue mechanics after depleting this protein extremely rapidly. The speed of depletion is important to avoid non-specific downstream effects, including possibilities ranging from compensatory mechanisms to severe cellular dysfunction. As a result, techniques such as gene mutation or RNAi are likely to be too slow-acting, while protein-based depletion techniques such as induced degradation or sequestration appear more promising (Natsume and Kanemaki, 2017).

A major debate in the field has emerged over whether embryonic tissues are predominantly viscous (Behrndt et al., 2012, Mayer et al., 2010, Munster et al., 2019) or elastic (Doubrovinski et al., 2017, Hernandez-Vega et al., 2017, Vuong-Brender et al., 2017) on the developmentally relevant timescale. A number of recent papers (by ourselves and others) demonstrated that after various embryonic tissues are stretched, they recoil back to the original configuration, suggesting they are elastic (Bambardekar et al., 2015, D’Angelo et al., 2019, Doubrovinski et al., 2017). However, those observations are consistent with viscosity-dominated models if the recoil is driven by active (notably myosin-generated) force. Distinguishing between these possibilities would require doing the measurements after molecular motors are depleted.

In the first hours of development, the fly embryo is a syncytium in which nuclei share a common cytoplasm as they undergo 13 rounds of division. Subsequently, in the process of cellularization, membranes protrude from the surface of the embryo in between those nuclei to partition the embryo into separate cells, thus forming a stage called the blastula (Campos-Ortega and Hartenstein, 1985). Immediately afterward, the large-scale tissue movements of gastrulation begin. These processes in *Drosophila* have served as a prime model for understanding the molecular and physical mechanisms underlying epithelial morphogenesis (Blankenship et al., 2006, Kiehart et al., 2017, Martin et al., 2009). Additionally, several techniques have been developed to examine the mechanical properties of the early embryo (Bambardekar et al., 2015, D’Angelo et al., 2019, Doubrovinski et al., 2017), making this an ideal system in which to investigate the contribution of individual cytoskeletal components to tissue rheology.

In this study, we used two complementary techniques to acutely inactivate myosin regulatory light chain (MRLC, called spaghetti squash or sqh in *Drosophila*) in the early Drosophila embryo. One of these techniques relies on the recently introduced AID2 system and was optimized to enable depletion after tens of minutes following injection of a drug. The second technique, introduced here, relies on conditional aggregation of a GFP-tagged protein, and operates within minutes. Combining these techniques with our previously developed method to probe material properties of embryonic tissues, we have established that mechanical properties of the early embryo are largely unaffected by sqh depletion. In combination with our previous findings, these results indicate that tissue recoil upon force removal is indeed a signature of elasticity rather than active tension.

## Results

To rapidly deplete the myosin regulatory light chain *sqh* in the early *Drosophila* embryo, we first used a recently enhanced version of the AID degron system (Nishimura et al., 2009, Yesbolatova et al., 2020). This technique involves translationally fusing the protein of interest to AID (auxin-inducible degron), which binds a plant-derived protein TIR1 (transport inhibitor response 1) upon addition of auxin. Once the AID-TIR1 complex is formed, the AID tag is polyubiquitinated and the AID-fusion protein targeted for proteasomal degradation. The AID2 system, which we use here, eliminates leaky (auxin-independent) degradation by employing a mutated TIR1 (TIR1^F74G^) and a modified ligand (5-Ph-IAA or 5-Ph-IAA-AM) (Negishi et al., 2022, Yesbolatova et al., 2020).

We first engineered a fly stain where *sqh* was translationally fused to a shortened AID sequence and GFP at the endogenous locus using CRISPR/Cas9 (Kubota et al., 2013, Morawska and Ulrich, 2013, Bence et al., 2017, Brosh et al., 2016). To decrease the total amount of myosin to be depleted, and thus the expected depletion time, we constructed flies in which the second allele of sqh was a null (sqh^AX3^). We also overexpressed TIR1^F74G^ maternally using two UASp-TIR1^F74G^ constructs driven by two copies of mat-GAL4 to further enhance the rate of myosin degradation upon drug application. Finally, the genotype used included a red membrane marker (Gap43-mCherry) to visualize cell outlines.

With this genetic background prepared, we began experiments investigating how myosin activity impacts tissue rheology in the early fly embryo. First, we examined the time-course of myosin depletion upon induction of AID-mediated degradation. Embryos were injected with 5-Ph-IAA-AM approximately 10-15 minutes prior to the onset of cellularization, then imaged to monitor the green fluorescence signal from any remaining sqh^GFP-AID^. Over multiple experiments, average fluorescence was found to drop to its minimum within 30-50 min after the injection (Figures 1a, 1c-c’). This minimum level of fluorescence is comparable to autofluorescence, as assessed by imaging embryos expressing Gap43-mCherry but not GFP, indicating that the pool of sqh is overwhelmingly depleted. Images at the level of the basal cellularization front (Figure 1b-b’) show that very faint fluorescence remains detectable in injected embryos at this location. However, basal outlines become markedly wavy and irregular (Figure 1b-b’ insets). This indicates that residual amounts of sqh^GFP-AID^ are insufficient to maintain basal tension.

**Figure 1.**
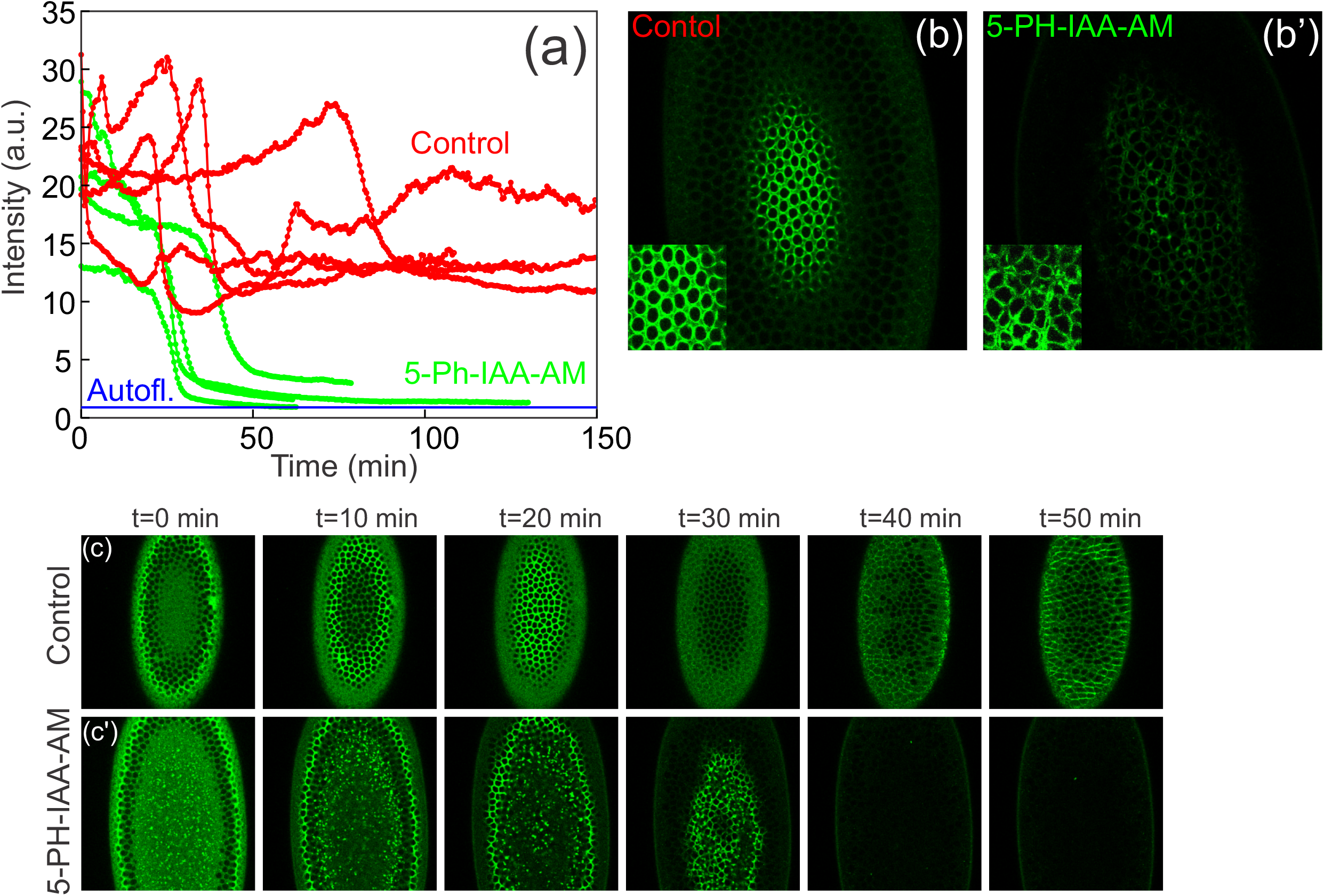
The kinetics of myosin depletion using the AID2 degron system. a). The dynamics of myosin depletion quantified as the time-course of fluorescence intensity within a region of interest. b-b’). Myosin fluorescence at the cellularization front in a control embryo and in a 5-Ph-IAA-AM injected embryo at the same time-point in cellularization. Confocal slices were taken 26 microns beneath the surface of the embryo. The main images were taken with the same microscope settings and processed in identical ways. The insets are magnified images, adjusted separately for brightness to allow for easier visualization in the 5-Ph-IAA-AM sample. c-c’). Representative snapshots showing the time-course of sqh^GFP-AID^ depletion.

Having established a method to rapidly deplete sqh in the early embryo, we asked how this influences material properties of embryonic tissues. To this end, we exploited a previously developed assay for probing tissue rheology using ferrofluid droplets (Doubrovinski et al., 2017). In this assay, a small (10-20 micron) droplet of ferrofluid is first injected into an embryo and positioned in the cellular layer by an externally applied magnet (Figure 2a). The magnet is then rapidly placed near the embryo surface some distance from the ferrofluid droplet, causing the droplet to move towards the magnet and stretching the surrounding cellular layer. In our experiments, we imposed this force-loading phase for approximately one minute (Figure 2a’). The magnet was then rapidly withdrawn, removing the external magnetic force from the droplet and allowing any subsequent tissue relaxation to be monitored. (Figure 2a’’).

**Figure 2.**
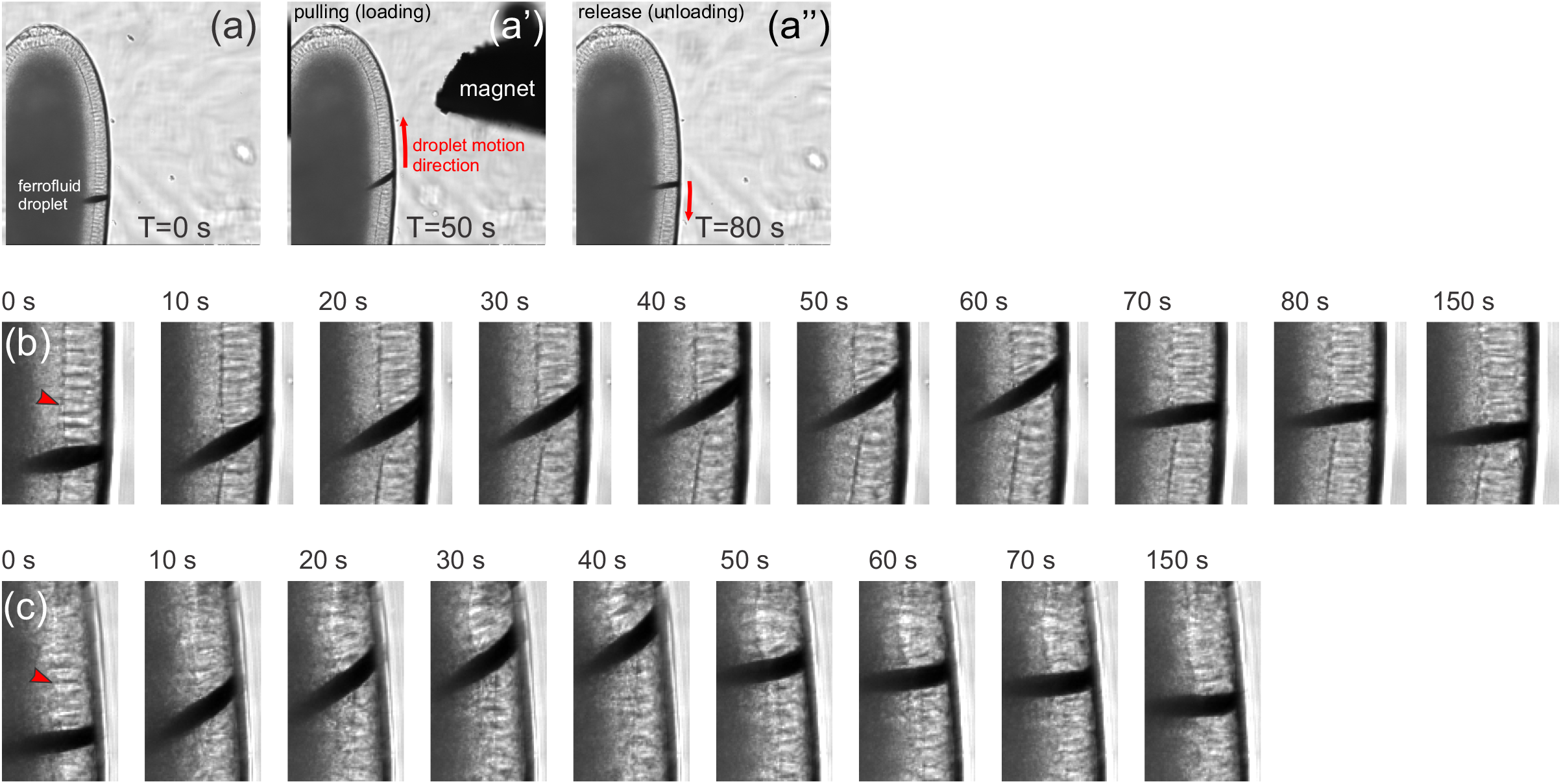
Ferrofluid-based assay used to quantify tissue material properties in different conditions. a-a’’). Snapshots of the ferrofluid pulling assay performed on untreated (control) embryos. b-c). Zoomed in regions of interest illustrating the dynamics of the droplet being pulled by the magnet in two experiments. b) is an uninjected control embryo, while c) is a 5-Ph-IAA-AM injected sqh^GFP-AID^ embryo. Note that the cellularization front (red arrowhead) is much more apparent in the untreated control case: this is a representative effect of the treatment, making the cellularization front appear “floppy” and uneven.

We applied this ferrofluid assay multiple times to both control embryos (Figure 2b) and embryos in which sqh was depleted using the AID2 system (Figure 2c). To allow for maximal sqh depletion, ferrofluid pulling experiments were performed 40 to 60 minutes after the injection of 5-Ph-IAA-AM. Qualitatively, the observed mechanical responses are essentially indistinguishable. In particular, during the force-loading phase the displacement of the droplet *d* as a function of time is well described by a power law dependence *d*(*t*) = *t^α^*, with *α* ≈ 1/2, in both the drug-treated embryos as well as in the controls. These results are in close quantitative agreement with previously reported data (Doubrovinski et al., 2017), indicating that the particular genetic background does not significantly affect tissue material properties.

To compare the two conditions more quantitatively, we calculated characteristic properties that quantify the dynamics in the unloading, or relaxation, phase of the assay. One characteristic property is the percent recoil, which reflects the extent to which elastic energy is stored over time. In other words, a system that stores elastic energy perfectly would have 100% recoil, while lower percentages indicate that the system “forgets” its initial state over time. For control experiments, the percent recoil measured 73.5±2.1, while the corresponding figure for the sqh-depleted embryos was 74.8±2.6, indicating that removal of sqh had no apparent effect on the ability of the tissue to store elastic stress. The other characteristic property we measured was the relaxation time constant (measured as the asymptotic slope of the decay curves in the rightmost plots from Figure 3), which reflects the ratio of elasticity to viscosity of the tissue. The relaxation time-constant was 64.5±5.0 seconds in the control condition, and 52.8±5.6 sec for the sqh-depleted embryos. The slight apparent difference is these time constants was not statistically significant (see Figure 3 legend); more importantly, though, it was remarkably small in magnitude. Together these results seem to indicate that the mechanical properties of the tissue are largely independent of sqh.

**Figure 3.**
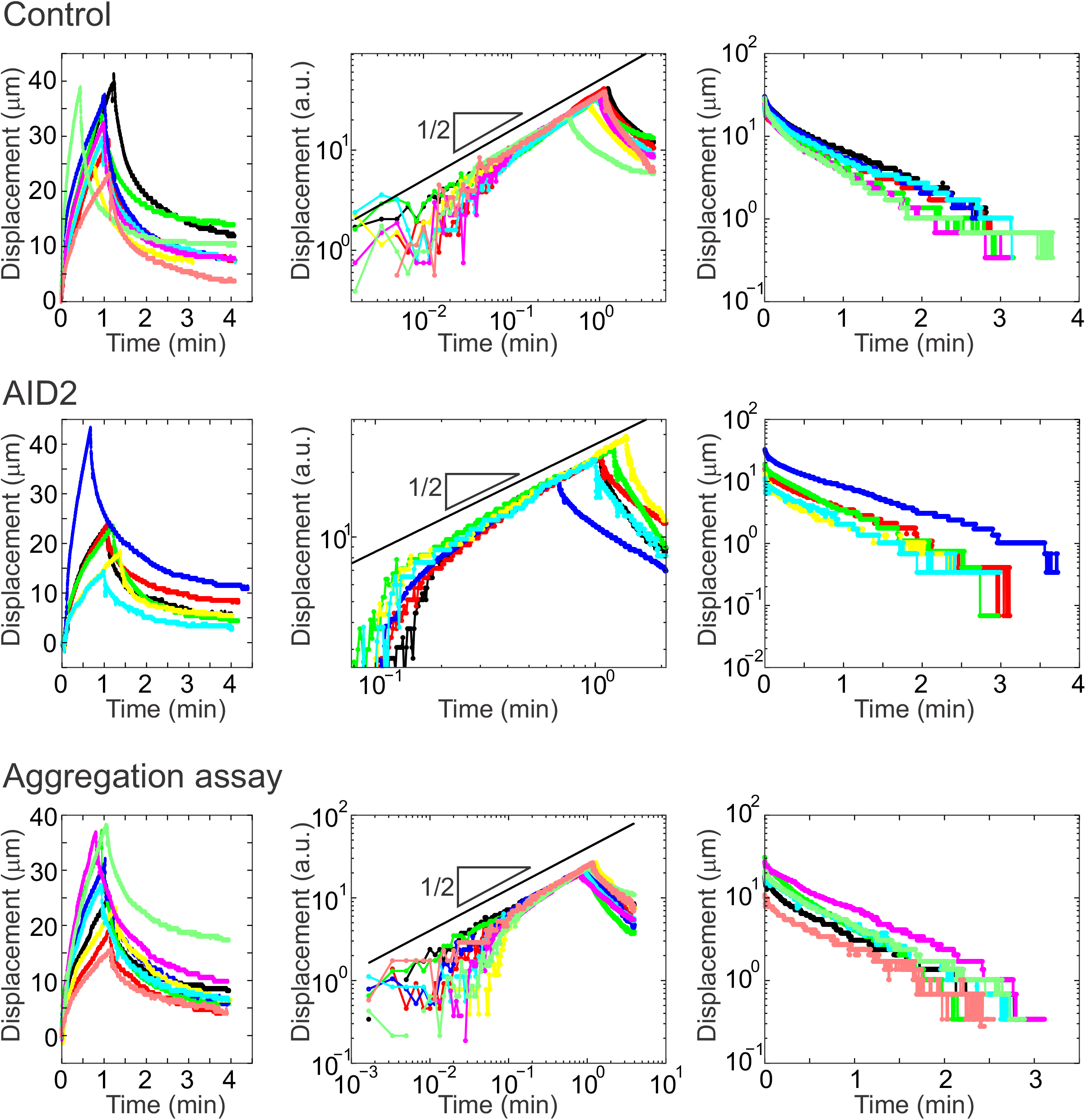
Quantification of tissue material properties after myosin depletion. Top panels show the measurements in the untreated (control) embryos. Middle panels correspond to embryos in which sqh^GFP-AID^ was depleted using the AID2 technique. Bottom panels correspond to embryos in which sqh^FRB-GFP^ was depleted using the nanobody-GFP aggregation-based technique. Left-most figures show the displacement of the ferrofluid droplet as a function of time. Middle figures represent the same data on a log-log plot. Right-most figures show plots of the recoil dynamics with logarithmic y-axis. Average values ± Standard Error of the Mean are given in the main text. Single-factor Anova failed to show a statistical difference among all three samples for the calculated relaxation time-constant (p=0.11) and percent recoil (p=0.92). Control n=9, AID2 n=6, Aggregation assay n=9.

Although our results from the AID2 degron assay suggest that tissue material properties are not substantially affected by depletion of sqh, we were concerned that the 40 minute interval allowed for protein degradation was relatively long and could still allow for downstream compensatory processes to occur. To address this concern, we sought to develop a technique to interfere with sqh on a still faster timescale. We decided to try sequestering our protein into non-functional aggregates. Our first attempt involved endogenously tagging sqh with FRB and GFP, and monitoring the aggregation of sqh upon injection of FKBP and rapamycin. Although the simple combination of FKBP and rapamycin injected together did not promote significant aggregation of sqh^FRB-GFP^ protein (data not shown), the injection of rapamycin and biotinylated FKBP followed by streptavidin resulted in rapid and pervasive aggregation of sqh^FRB-GFP^ in the vicinity of the injection site. Furthermore, this aggregation appeared to deplete sqh activity, since cellularization was markedly slowed and basal rings failed to close in this region (Figure 4c,c’).

**Figure 4.**
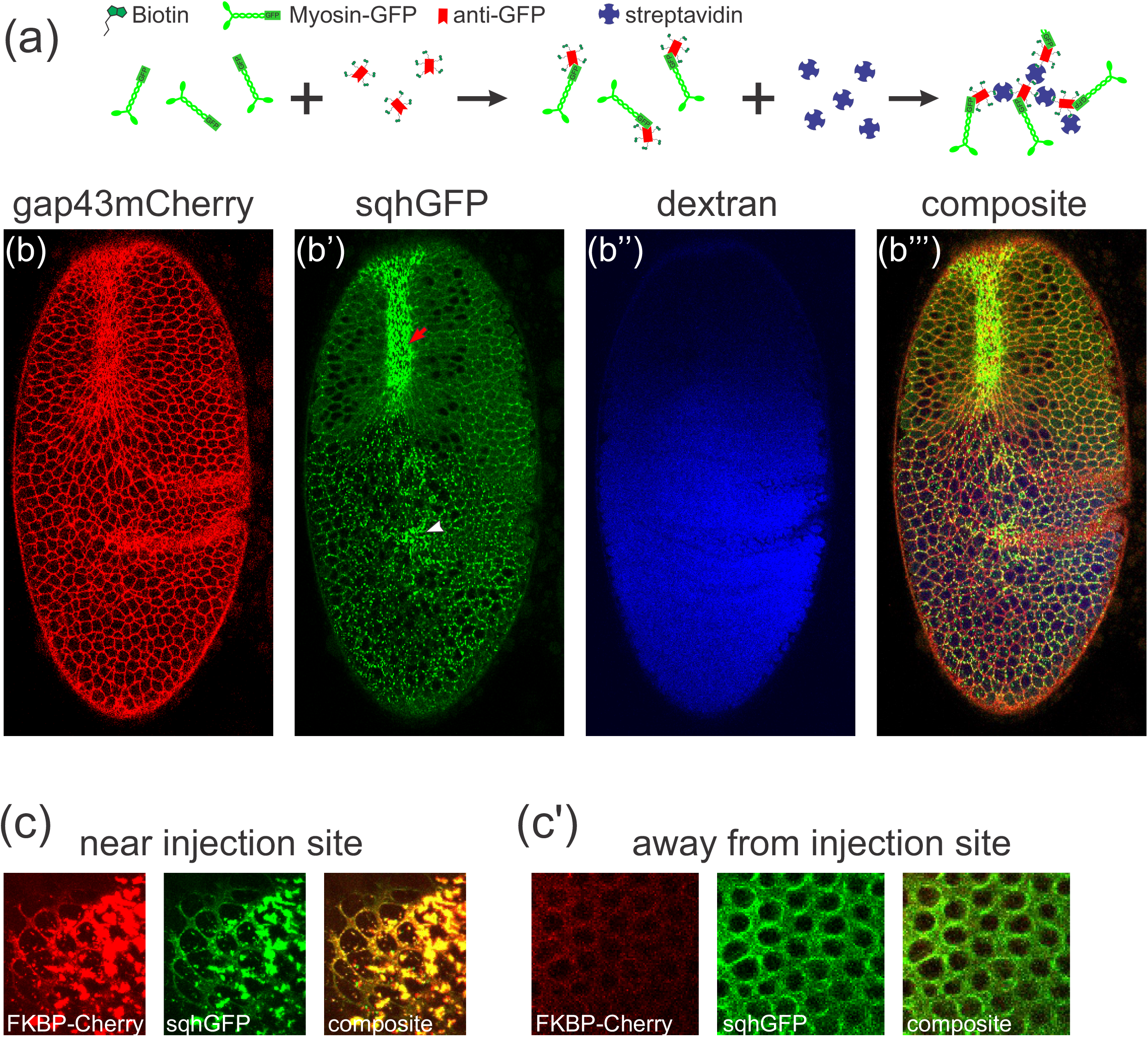
Aggregation-based assay for depletion of abundant proteins in the early *Drosophila* embryo. a) Schematic illustrating the basis of the method (see text). b-b’’’) A representative sqh^FRB-GFP^ embryo that has been injected with biotinylated anti-GFP nanobody and streptavidin. Blue signal (b’’) is co-injected dextran indicating distribution of the injected material. Gastrulation fails in the vicinity of the injection site. Red arrow indicates region with properly forming ventral furrow. White arrowhead indicates region where sqh^FRB-GFP^ signal forms aggregates; the ventral furrow fails to form in this region. c-c’). A representative sqh^FRB-GFP^ embryo that has been injected with rapamycin, biotinylated FKBP, and streptavidin. All images were taken from the same z-position and time point, showing that basal rings fail to close in the vicinity of the injection site.

Inspired by these results, we next attempted to develop a more generally applicable aggregation technique using several layers of multivalent interactions, but targeting the GFP tag rather than the FRB tag (Figure 4a). We found that injecting sqh^FRB-GFP^ embryos with biotinylated anti-GFP nanobody followed by streptavidin resulted in as rapid and extensive aggregation of sqh^FRB-GFP^ as seen with our FRB-based technique (Figure 4b-b’’’). This approach also resulted in the apparent loss of sqh activity in the injected region, as seen by defects in ventral furrow formation (Figure 4b-b’’’). This approach is also remarkably fast compared to AID2-based degradation; sqh^FRB-GFP^ aggregation appears to occur completely within the approximately 5 minutes it takes to move the sample from our injection set-up to the confocal microscope for imaging. This allows us to examine the effects of sqh on tissue rheology while further reducing the possible action of compensatory downstream effects. We then performed ferrofluid-based measurements on embryos where myosin was depleted through aggregation and found that tissue mechanics again remained largely unaffected (Figure 3). Specifically, the percent recoil measured 73.4±2.8, while the relaxation time-constant of the recoil had a value of 50.6±3.5 sec. In this way, this much more rapid technique for protein depletion corroborates our previous finding using the AID2 system.

## Discussion

In this study, we used the AID2 degron technique and a novel sequestration approach to acutely inactivate the myosin regulatory light chain sqh in the early Drosophila embryo and examine the effect on tissue material properties. We demonstrated that both techniques were very effective at depleting myosin activity, as seen by the resulting cellularization and ventral formation phenotypes. Strikingly, sqh depletion causes little to no change in tissue rheology. Specifically, we compared measurements reflecting the storage of elastic energy over time and the ratio of elasticity to viscosity, and found that neither was significantly altered.

Several studies have explored cell and tissue mechanics by measuring the deformations induced under some particular pattern of applied force (Bambardekar et al., 2015, Bausch et al., 1998, D’Angelo et al., 2019, Doubrovinski et al., 2017, Fernandez et al., 2007, Hoffman et al., 2006, Khalilgharibi et al., 2019, Thoumine and Ott, 1997). However, the interpretation of these measurements is not always straightforward (Wu et al., 2018). One complication comes in trying disentangle the effects of passive properties versus active (i.e. energy-consuming) responses. Importantly, a meshwork that exhibits very low elasticity but is subjected to significant uniform active stress can exhibit mechanical responses very similar to a network with high elasticity and no stress present (Mulla et al., 2019). In previous work in cellularization-stage Drosophila embryos, we found that tissue recoil after removal of force is nearly abolished in the presence of cytochalasin D, and we argued that the recoil should be interpreted as the passive elastic response of the actin cytoskeleton (Doubrovinski et al., 2017). However, an alternative interpretation exists: the recoil is an active process due to myosin contractility. The results of the current study argue strongly against this alternative explanation, and support the conclusion that the epithelium of cellularization-stage Drosophila embryo is highly elastic.

Beyond supporting the conclusion that this tissue exhibits strong passive elasticity, the results presented here also indicate that myosin activity has little to no influence on the strength of that elasticity. This is an important thing to establish in our system of interest, since experiments in other cells and tissues have variously shown stiffening (Chan et al., 2015), softening (Martens and Radmacher, 2008), or no effect on elasticity (Fernandez et al., 2007, Van Citters et al., 2006) in response to myosin inhibition.

Since the experiments here involved specifically depleting the myosin regulatory light chain, we were probing the contribution of active myosin. This may not be identical to the contribution of the myosin complex overall. Work in other systems has suggested that the myosin heavy chain alone can act as an actin crosslinker that makes substantial contributions to elasticity (Xu et al., 2001). In the future, it would be interesting to test whether myosin heavy chain acts similarly in this tissue, by targeting the non-muscle myosin 2 heavy chain zipper for degradation or sequestration and measuring the impact on rheology.

On the technical side, we have introduced two related novel techniques (based on FKBP-FRB and nanobody-GFP interactions) for very rapid sequestration of highly abundant proteins in the early embryo. Since GFP tags have been introduced into a large number of genes in the Drosophila genome (Buszczak et al., 2007, Morin et al., 2001, Quinones-Coello et al., 2007, Nagarkar-Jaiswal et al., 2015), the nanobody-GFP based technique is immediately applicable to studying the role of those proteins in the embryo. Both techniques involve the injection of some components, which limits their application, but are still accessible in a variety of key models systems such as tissue culture cells, oocytes, C. elegans germline, and other syncytial tissues. Although several techniques based on similar principles have been developed recently (Hernandez-Candia et al., 2021, Lee et al., 2014, Qin et al., 2017, Yoshikawa et al., 2021), both the rate and the degree of protein inactivation achieved by our method is significantly higher than in the previous studies. We anticipate that many of the proteins that control tissue mechanical properties may be structural, rather than regulatory, in nature and therefore present in high concentration. A highly efficient method of depletion is therefore key to understanding how such proteins contribute to tissue mechanics. More generally, we envision that our approach of combining acute protein degradation or sequestration with rheological measurements will find additional applications in many other systems, and aid in dissecting the contributions of individual proteins to material properties in living biological systems.

## Materials and Methods

### Plasmid construction

The gap43_mCherry_pCaSpeR4 plasmid was generated by inserting a sequence encoding the gap43 tag (the first 20 amino acids of rat gap43) translationally fused to mCherry between the NotI and XbaI sites of Tubp-Gal80 in pCaSpeR4 (a gift from Liqun Luo; Addgene plasmid # 17748; http://n2t.net/addgene:17748; RRID:Addgene_17748) (Lee and Luo, 1999). This eliminated the Gal80 sequence and resulted in a plasmid with gap43-mCherry downstream of the tubulin promoter.

The mat-Gal4-sqhUTR plasmid was generated by PCR amplification of the Gal4 coding sequence from pPac-Pl-mCD8-D2A-Gal4 (a gift from Benjamin White, Addgene plasmid # 39457; http://n2t.net/addgene:39457; RRID:Addgene_39457) (Diao and White, 2012). Primers used included a 5’ NheI site and Kozak sequence (ATCAAC) and a 3’ NotI site. The PCR product was inserted it into NheI/NotI cut alphaTub67c_MCS_sqhUTR pBABR (a gift from Shelby Blythe).

The pUASP-attB-TIR2-F74G plasmid was generated by synthesizing the TIR1-F74G sequence, along with a 5’ Kozak sequence and 3’ linker sequence (VDSGSAASG) plus HA tag, and inserting this into UASp-attB (DGRC Stock 1358; https://dgrc.bio.indiana.edu//stock/1358; RRID:DGRC_1358) between NotI and XbaI. The TIR1-F74G sequence was a gift from Masato Kanemaki (Addgene plasmid # 140730; http://n2t.net/addgene:140730; RRID:Addgene_140730) (Yesbolatova et al., 2020).

The sqh_GFP_FRB_donor plasmid was generated by inserting synthesized DNA between the two EcoRI sites of ScarlessDsRed_pHD-2xHA (a gift from Kate O’Connor-Giles, Addgene plasmid # 80822; http://n2t.net/addgene:80822; RRID:Addgene_80822). The inserted DNA contained these elements in order: a 900 base pair homology arm derived from the sqh genomic sequence and ending with the final sqh coding exon; a linker encoding DYKEPVAT; GFP(S65T); FRB; a stop codon; and a 900 base pair homology arm corresponding the genomic sequence downstream of the sqh coding region.

The sqh-GFPaid_v2 plasmid was also generated by inserting synthesized DNA into the ScarlessDsRed_pHD-2xHA backbone between the two EcoRI sites. The inserted DNA is the same as in the sqh_GFP_FRB_donor plasmid with two exceptions: the sequence encoding AID (from Genbank NM_100306.4) was substituted for FRB, and the first homology arm including a single, silent A-to-G change ten bases before its end to mutate the PAM site targeted by one of the guide RNAs.

The two guide RNA-expressing plasmids used for modification of the sqh genomic locus were designed using the tool at https://flycrisper.org/target-finder (Gratz et al., 2014). The following sense/antisense pairs oligos were annealed separately and ligated into the BbsI sites of pU6-BbsI-chiRNA (Gratz et al., 2013) 194:1029): guide1-sense: CTTCGCTGGCTAGTGAATCGACTGC; guide1-antisense: AAACGCAGTCGATTCACTAGCCAGC; guide2-sense: CTTCGTAAGCACGGTGCCAAGGACA; guide2-antisense: AAACTGTCCTTGGCACCGTGCTTAC.

pET_FRB_Dark Cherry_1xNano encodes a fusion protein containing 6xHis tag, dark mCherry, and anti-GFP nanobody sequences. The dark mCherry sequence was created by making a Y-to-G mutation in the chromophore. The GFP-Nanobody sequence was a gift from Kazuhisa Nakayama (Addgene plasmid # 61838; http://n2t.net/addgene:61838; RRID:Addgene_61838) (Katoh et al., 2015). pET_FKBP_mCherry encodes a fusion protein containing 6xHis tag, FRB, and mCherry sequences. pET_FRB_Dark Cherry_1xNano and pET_FKBP_mCherry plasmids were constructed from pET28 FRB-ferritin and pET28 FKBP-ferritin respectively (both kind gifts of Zoher Gueroui) (Ducasse et al., 2017). The complete annotated sequences are listed in the supplement.

### Fly stocks

The genotype used for AID-degron experiments was sqh^GFP-AID^, gap43-mCherry/sqh^AX3^; +; mat-GAL4^89E11^, mat-GAL4^15^/UASp-TIR1-F74G^VK00027^, UASp-TIR1-F74G^attP2^. For aggregation experiments, the genotype used was sqh^FRB-GFP^;+; gap43mCherry/TM3, Sb. The control line used to measure background green autofluorescence in Figure 1A was w;+; gap43mCherry/TM3, Sb.

Bloomington Drosophila Stock Center was the source of mat-GAL4^15^ (RRID:BDSC_80361) and sqh^AX3^ (RRID:BDSC_25712). The gap43-mCherry (membrane-mCherry) on the third chromosome was a gift from Adam Martin (Martin et al., 2010).

Flies with gap43-mCherry on the first chromosome were created by injection of the gap43_mCherry_pCaSpeR4 plasmid into fly embryos using P-element mediated insertion; progeny were screened for X chromosome transformants. mat-GAL4^89E11^ flies were created by injection of the mat-Gal4-sqhUTR plasmid into Rainbow line 35569 (genotype: y^1^ w[*] P{ y[+t7.7]=nos-phiC31int.NLS} X;+; PBac{ y[+]-attP-9A} VK00027). UASp-TIR1-F74G^VK00027^ and UASp-TIR1-F74G^attP2^ flies were created by injection of the pUASP-attB-TIR2-F74G plasmid into the Rainbow lines 35569 (genotype listed above) and R8622 (genotype: M{vas-int.Dm}ZH-2A;+; P{CaryP} attP2), respectively.

Flies carrying sqh^FRB-GFP^ and sqh^GFP-AID^ were generated by CRISPR-based modification to the C-terminus of *sqh* at its endogenous locus. In both cases, flies of genotype y,sc,v; +/+; nos-Cas9 attP2 were injected with a mixture of 250 ng/ul donor plasmid and 20 ng/ul of each guide plasmid in water. The same two guide plasmids (described in the Plasmids section) were used in both cases. The donor plasmids for sqh^FRB-GFP^ and sqh^GFP-AID^, called sqh_GFP_FRB_donor and sqh-GFPaid_v2 respectively, are also described in that section. Progeny of injected flies were screened for successful modification of the sqh genomic locus by a combination of screening for GFP fluorescence in dissected egg chambers and genomic PCR with sequencing. The sqh^FRB-GFP^ flies generated by this method incorporated the full length FRB-GFP sequence from the donor plasmid. Only one sqh^GFP-AID^ line was found screening by GFP fluorescence, and genomic sequencing revealed that while the entire GFP sequence was incorporated, the AID sequence incorporated was truncated, followed by a small inserted sequence, then the 3’ UTR of sqh. The final result is a shortened AID tag (M1-K124) followed by the amino acid sequence DEQ. Since this shortened tag included a functional shortened mini-AID sequence (Brosh et al., 2016, Kubota et al., 2013, Morawska and Ulrich, 2013) (Kubota et al 2013, Morawska and Ulrich 2013, Brosh et al 2016), we tested and successfully used this line in experiments.

All injections for generating transgenic flies were done by Rainbow Transgenic Flies, Inc.

### Protein purification and biotinylation

Anti-GFP nanobody protein was produced from the plasmid pET_FRB_Dark Cherry_1xNano. The final fusion protein includes FRB (FKBP-rapamycin binding) and dark mCherry domains, but these do not play specific functional roles in the experiments presented here (beyond providing more potential biotinylation sites), so we refer to this protein as “anti-GFP nanobody” for clarity. FKBP-mCherry protein (in Supplemental Figure S1) was produced from the plasmid pET_FKBP_mCherry.

To produce protein, expression plasmids were transformed into *E. coli* BL21(DE3) cells (ThermoFisher cat# C600003). Overnight cultures were used to inoculate flasks containing 500 mL lysogeny broth (LB) with 50 μg/mL kanamycin, and flasks were shaken at 37 °C until cultures reached an OD_600_ of 0.6–0.8. Isopropyl β-D-1-thiogalactopyranoside (IPTG) (Invitrogen #15529-019) was added at a final concentration of 1 mM and the cultures shaken at 26 °C for 3 hours. Cells were harvested by centrifugation and pellets were frozen at −80 °C for later use. Thawed cell lysates were resuspended in suspension buffer (100 mM NaCl, 4 mM Tris pH 7.5, supplemented with cOmplete, EDTA-free Protease inhibitor cocktail, Sigma cat # 11873580001). Cells were lysed at 4 °C by incubating with 0.002 microgram/ml lysozyme (Sigma L6876) for 10 minutes, sonicated for 4 10-second pulses in a MisonX 3000 ultrasonicator, and incubated with 0.05 mg/ml Deoxyribonuclease I (Sigma D4527), 12.5 micromolar MgCl_2_, and 1.25% Triton X-100. Lysate was cleared by centrifugation at 24,000*g* for 30 min.

Proteins were His-tagged, and so were purified using Ni-NTA agarose beads (ThermoFisher Scientific cat# R90101) essentially according to manufacturer’s instructions. Protein recovery was maximized by adjusting the concentration of imidazole (Sigma cat# l5513-5G) as follows. Ni-NTA agarose beads (ThermoFisher Scientific cat# R90101) were resuspended in 0 mM Imidizole buffer (50mM sodium phosphate; pH 8.0; 300mM NaCl) before incubating with cleared lysate. Beads were washed twice with 0 mM Imidizole buffer, then twice with 10mM Imidizole buffer (50 mM sodium phosphate pH 8.0; 300 mM NaCl; 10 mM imidazole). 500 microliter fractions were eluted with 250mM Imidizole buffer (50mM sodium phosphate pH 8.0; 300mM NaCl; 250 mM imidazole), and protein concentration was measured by OD_280_. Fractions of at least 1 mg/ml protein were combined and dialyzed into PBS using 10K MWCO dialysis cassettes. For the data presented in this paper, the concentration of FKBP-Cherry as measured before biotinylation was 12 mg/ml, and the concentration of anti-GFP nanobody as measured before biotinylation was 6.4 mg/ml, but a range of concentrations used in other experiments gave similar results.

To biotinylate the proteins, Biotinamidocaproate N-Hydroxysuccinimide ester (Sigma # B2643) was dissolved to 38.5 mM in N,N-dimethylformamide (Sigma-Aldrich #227056) then combined with the protein solution in a 15:1 (biotinylation reagent : protein) molar ratio. After incubation for 45 minutes at room temperature, the protein was dialyzed into PBS to remove unconjugated biotin and stored at 4 °C.

### Embryo Injections

Embryos were collected on grape agar plates and stage-selected under Halocarbon 27 oil (Sigma or Halocarbon InfinX), dechorionated in bleach, then washed with water. Embryos were then mounted ventral side down along a line of embryo glue (double-sided scotch tape extract in heptane) on a glass coverslip. The coverslip was placed in a closed plastic container with drierite, and allowed to dry for 15 minutes. Embryos were then covered in Halocarbon 700 oil (Sigma) to prevent further drying. Embryos were injected laterally, near the anterior end.

For AID degron experiments, embryos were injected with 50 mM 5-Ph-IAA-AM in DMSO. For aggregation experiments, embryos were first injected with biotinylated anti-GFP nanobody (or biotinylated FKBP-mCherry mixed with equimolar rapamycin, centrifuged to clear, in Supplemental Figure S1) solution on one side of the embryo. The coverslip slip was then rotated 180 degrees, and a streptavidin mixture was injected laterally, directly opposite the first injection. This mixture was prepared by mixing 20 mg/ml streptavidin (Roche 11721674001) in a 2:1 ratio with 10 mg/ml Alexa Fluor 647 conjugated dextran (ThermoFisher D22914), to aid in visualizing injection into the embryo. Ferrofluid injections, if any, were performed last. All injections were facilitated using a FemptoJet 5247 Microinjector (Eppendorf).

To prepare needles for protein or 5-Ph-IAA injection, glass capillaries (Sutter Instruments, #BF100-78-10) were pulled using a micropipette puller (Sutter Model P-2000) using the following settings: Heat=350, Fil=6, Vel=30, Del=110, Pul=120. Needles were loaded using microloader tips (Eppendorf 5242 956.003). Finally, the needle was gently broken by pressing the tip against the side of a coverslip after loading to open the tip.

### Ferrofluid experiments

Ferrofluid injections and rheological measurements were done as previously described in (Doubrovinski et al. 2017). Borosilicate glass capillaries with an inner diameter of 0.5 mm (Sutter BF100-50-10) were pulled into injection needles using a Sutter P-2000 pipette puller with the following settings: Heat=380, Fil=4, Vel=20, Del=128, Pul=120. Pulled needles were beveled using a EG-45 microgrinder (Narishige), then loaded with ferrofluid (Apexmagnets Ferrofluid Magnetic Liquid - 8 oz) for injection into embryos. External force was applied to ferrofluid droplets using a permanent magnet mounted on a manual translation stage.

### Imaging

Imaging was done on a Zeiss LSM 700. A Plan-Apochromat 20x/0.8 M27 objective was used for ferrofluid experiments; an EC Plan-Neofluar 40x/1.30 oil DIC M27 objective was used for all other experiments.

## Acknowledgements

Plasmids obtained from the Drosophila Genomics Resource Center (NIH Grant 2P40OD010949) and stocks obtained from the Bloomington Drosophila Stock Center (NIH P400D018537) were used in this study.

